# reserBUGS: A reservoir computing framework for probabilistic forecasting of ecological abundance time series

**DOI:** 10.64898/2026.08.14.744603

**Authors:** Miguel A. Mohedano-Muñoz, Javier Galeano, Juan M. Pastor, Julia G. de Aledo, Ignasi Bartomeus, Alfonso Allen-Perkins

## Abstract

Forecasting species population dynamics is a central challenge in computational ecology, yet existing approaches rarely combine flexible nonlinear modelling, support for count-based ecological data, and systematic uncertainty quantification within a single, scalable framework. Here we introduce `reserBUGS`, an open-source Python framework for ecological forecasting based on reservoir computing, a recurrent neural network architecture in which only a simple readout layer is trained while a fixed high-dimensional dynamical system encodes temporal memory and nonlinear dependencies. `reserBUGS` integrates species abundance time series with environmental covariates retrieved automatically from global climate products, generates probabilistic ensemble forecasts, and provides tools for forecast evaluation and reliability assessment. We evaluated `reserBUGS` using insect abundance time series from available biodiversity monitoring datasets, comparing its performance against seven statistical and machine-learning baselines over one- to five-year forecast horizons. Reservoir-based models consistently outperformed alternatives in both predicting future abundance and capturing forecast uncertainty, with environmental predictors increasing the proportion of stable forecasts and contributing additional predictive value beyond historical abundance dynamics alone, particularly at 3-4-year forecast horizons. Probabilistic forecasts further enabled the identification of conditions associated with reduced predictive skill, providing a practical basis for communicating forecast confidence to end users. While default configurations already achieved competitive performance across a taxonomically and geographically diverse set of time series, hyperparameter optimisation revealed substantial room for performance gains through series-specific tuning. `reserBUGS` offers a computationally efficient and extensible framework for ecological forecasting that is well suited to the short, heterogeneous time series typical of biodiversity monitoring programmes. Its combination of flexible nonlinear modelling, probabilistic uncertainty quantification, and automated environmental data integration addresses key practical barriers to the adoption of modern forecasting methods in conservation and ecological research.

**Author summary:** Predicting how animal populations will change in the future is crucial for conservation, yet it remains surprisingly difficult. Population sizes fluctuate due to a complex mix of factors — climate, habitat, species interactions, and chance — making future dynamics hard to anticipate, especially when the available data are limited. We present `reserBUGS`, a computational tool designed to forecast animal population abundance. The tool is built around a technique called reservoir computing, which uses a large network of randomly connected artificial neurons to capture complex temporal patterns in data. We tested `reserBUGS` on thousands of insect population time series collected worldwide, and covering hundreds of species from butterflies to beetles. We found that our approach predicted future population sizes more accurately than commonly used statistical and machine-learning alternatives. Incorporating environmental information — such as temperature and precipitation — further improved forecast accuracy, particularly for predictions several years into the future. By providing estimates of forecast uncertainty, users can identify situations where predictions are likely to be less accurate. `reserBUGS` will be a useful tool for ecologists and conservation practitioners who need to anticipate population trends and make informed management decisions under uncertainty.

## Introduction

Ecological systems are undergoing human-induced rapid environmental changes. This sustained exposure to pressures such as climate change and land-use transformation is triggering profound changes in species abundances and diversity [1, 2]. Against the risk of ecologists becoming simple accountants of biodiversity loss, there is an increasing urgency to use conservation tools to reverse these trends. This implies that we need to better anticipate future biodiversity dynamics to act efficiently. Hence, ecology has increasingly embraced forecasting as a core scientific goal, complementing traditional explanatory and retrospective analyses [3]. However, prediction is particularly challenging for complex systems such as ecological communities [4, 5] and remains an open challenge for ecologists. As biodiversity monitoring expands in spatial extent and temporal resolution, the development of robust, scalable, and transparent forecasting methods becomes a central priority for computational ecology [3].

Forecasting ecological dynamics is challenging because ecological time series often exhibit properties that complicate the application of many standard modeling approaches [3, 6]. Species abundance time series, one of the data cornerstones for assessing species conservation status and trends [7], are typically short, noisy, and nonstationary [8–10], reflecting both intrinsic population processes and extrinsic environmental forcing [6]. Moreover, ecological responses to drivers such as climate, vegetation, or resource availability frequently occur with unknown and nonlinear time lags [11], generating dynamics that depend on extended histories rather than recent conditions alone [6]. Together, these features make it difficult to disentangle signal from noise and require forecasting methods that can flexibly accommodate memory, nonlinearity, and uncertainty.

A wide range of approaches has been applied to ecological forecasting, ranging from classical statistical frameworks, such as autoregressive and state-space models, to modern machine-learning approaches, including deep-learning architectures and other data-driven techniques [12–14]. At the core, these two approaches differ in the balance they strike between model interpretability and predictive flexibility, the latter often relying on black box models. On one end, autoregressive and state-space models [15] provide interpretable frameworks and often perform well under relatively simple dynamics, but may require careful specification and sufficient data to capture strong nonlinearities, non-stationarity, and long-range temporal dependencies [16]. At the other extreme, deep learning architectures such as recurrent neural networks and transformers offer substantial flexibility, but often require large training datasets that are rarely available in ecological monitoring programs, as well as extensive tuning and considerable computational resources [12, 14]. These trade-offs can limit how well different approaches perform on the short and heterogeneous time series typical of ecological monitoring data. Most importantly, these trade-offs highlight a broader challenge: existing approaches rarely combine flexible nonlinear modeling, support for count-based ecological data, and systematic forecast evaluation within a single, easy-to-use framework.

As a result, there remains a gap between the strengths and weaknesses of modern time-series models and their practical use in routine ecological forecasting, particularly when predictions need to be scalable, repeatable, and regularly evaluated. This challenge is further compounded by intrinsic limits to ecological predictability, which depend on system dynamics, data availability, and evaluation criteria [17–19], as well as by the practical difficulties of calibrating and validating ecological models under realistic data constraints [20].

In this context, a practical forecasting framework for ecology would combine the flexibility and predictive power of machine learning with the data efficiency required to handle the short, heterogeneous time series typical of biodiversity monitoring. *Reservoir computing* offers a promising path toward this goal. In reservoir computing, a fixed, high-dimensional dynamical system transforms input time series into rich internal representations that encode temporal memory and non-linear dynamics, while only a simple readout layer is trained [21, 22]. This separation enables efficient learning, particularly relative to fully trained recurrent networks, and can help mitigate overfitting in data-limited settings [22]. Because past inputs persist in the reservoir’s internal state, reservoir computing can also capture history-dependent and delayed effects, which are common in ecological systems. Indeed, reservoir computing has demonstrated strong performance in forecasting complex dynamical systems, including chaotic dynamics as well as applications in physics and climate science [22–24]. However, its adoption in ecology has remained comparatively limited, in part due to the relative scarcity of tools tailored to ecological data structures. Bridging this gap requires translating reservoir computing into frameworks that accommodate ecological data constraints and forecasting applications.

Recent work has highlighted the potential of near-term ecological forecasting, in which models generate short-horizon, quantitative predictions that can be evaluated against subsequent observations [25]. This approach supports adaptive management and fosters cumulative learning across systems and datasets. This implies that effective forecasting requires more than a stand-alone model and needs integrated pipelines that link data acquisition, preprocessing, model fitting, forecast generation, and evaluation within a coherent workflow [3]. Existing ecological forecasting frameworks are often bespoke or optimized for specific case studies, limiting their scalability and reuse across systems [3, 26]. Consequently, there is a need for forecasting frameworks that integrate modeling approaches within transparent, reusable, and extensible workflows for biodiversity time series.

Here we introduce `reserBUGS`, a reservoir-computing-based forecasting framework for ecological abundance time series [27]. `reserBUGS` provides: (i) an implementation of reservoir computing tailored to short and heterogeneous ecological time series, (ii) an integrated workflow linking data retrieval, including abiotic and satellite-derived predictors, with model fitting, forecasting, and evaluation, and (iii) support for probabilistic forecasting with ecologically relevant performance metrics, including metrics for quantifying forecast magnitude error.

Crucially, we evaluate `reserBUGS` performance using 2,858 insect abundance time series from large-scale biodiversity monitoring datasets [28], and compare its forecasting performance against seven established statistical and machine-learning baselines, as well as a reservoir model driven solely by historical abundance. Insects play a central role in ecosystem functioning and the provision of ecosystem services such as pollination and pest control [29], while exhibiting substantial spatial and temporal variability in population trends across regions and taxa [10, 30]. These characteristics, together with their typically rapid life cycles [31, 32] and the relative scarcity of long-term standardized monitoring data [30, 33], make insect populations particularly challenging to forecast. Their dynamics are often difficult to capture using conventional modeling approaches [3, 6], providing a stringent test case for evaluating ecological forecasting methods.

We address three related questions. First, does reservoir computing improve the accuracy and probabilistic quality of ecological forecasts relative to established statistical and machine-learning approaches, and if so, to what extent does environmental information contribute to this improvement beyond that contained in historical abundance dynamics alone? Second, can probabilistic forecasts be used to quantify forecast uncertainty and identify conditions associated with reduced predictive skill? Finally, how sensitive is reservoir forecasting performance to hyperparameter selection? By addressing these questions, we evaluate the ability of `reserBUGS` to generate accurate probabilistic forecasts of ecological dynamics and to provide uncertainty estimates that inform confidence in ecological predictions.

## Materials and methods

### The reserBUGS forecasting framework

`reserBUGS` is an ecological forecasting framework that combines reservoir computing with environmental predictors to generate probabilistic forecasts of population abundance. The framework integrates ecological time series, environmental covariates, model fitting, ensemble forecasting, uncertainty quantification, and forecast evaluation within a common workflow (Figure 1).

**Fig 1.**
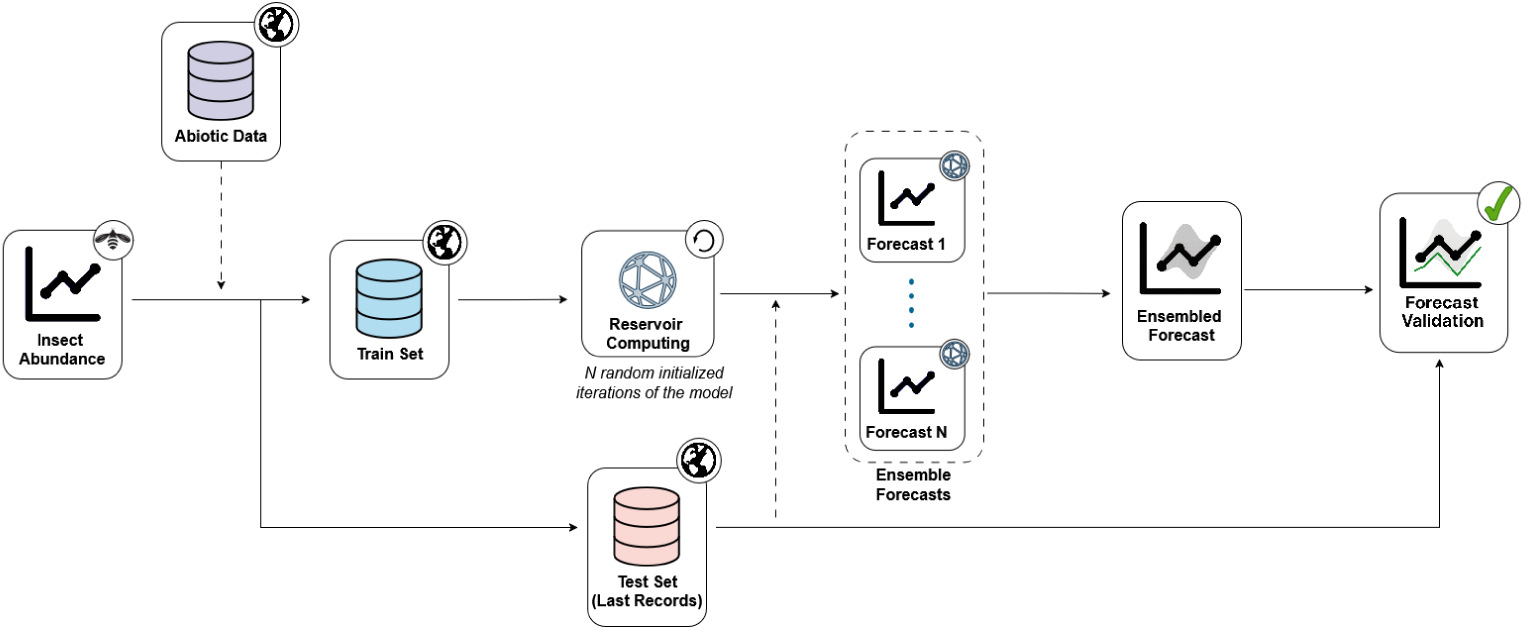
Overview of the `reserBUGS` ecological forecasting workflow. Species abundance time series derived from biodiversity monitoring data are combined with abiotic environmental predictors obtained from global climate reanalysis products. The resulting dataset is partitioned into training and test periods, and the training data are used to fit a reservoir-computing model. Multiple forecast trajectories are generated through repeated model realizations and combined into an ensemble forecast that provides both point predictions and predictive uncertainty. Forecast performance is subsequently evaluated against independent observations from the test period.

Reservoir computing [34] is a recurrent neural-network approach designed for modelling temporal dynamics. Rather than fitting a complex dynamical model directly, input variables are transformed into a high-dimensional representation of system dynamics, referred to as the reservoir, which retains information about recent system states and temporal dependencies. In this sense, reservoir computing shares similarities with gated recurrent architectures such as Long Short-Term Memory (LSTM) networks [35], as both aim to capture long-range temporal dependencies. However, unlike LSTMs, in reservoir computing only the output layer is trained, while the recurrent reservoir remains fixed, resulting in a simpler and computationally efficient training process [36]. The output layer transforms the reservoir states into predictions of the response variable.

In `reserBUGS`, this output layer is implemented as a generalized linear model (GLM). The framework supports Poisson, Gamma, and Tweedie GLMs, allowing the response distribution to be selected according to the characteristics of the data. In the present study, we use a Tweedie GLM because it provides a flexible framework for modelling non-negative, right-skewed abundance data that may include both exact zeros and positive values [37].

Reservoir dynamics are governed by a small set of hyperparameters, including reservoir size (i.e., the number of neurons in the reservoir), spectral radius (which determines the strength of recurrent interactions and thus influences memory), and leaking rate (which controls the rate at which neuron states are updated), which together determine the reservoir’s dimensionality and temporal memory [34].

Predictive uncertainty associated with the forecasting procedure was approximated through ensemble forecasting. For each time series, we generated *N* = 500 forecast trajectories by refitting independent reservoir realizations with different random seeds and with reservoir hyperparameters sampled from predefined uniform ranges. Specifically, spectral radius, leaking rate, and input scaling were sampled from the intervals 0.85–1.00, 0.75–0.95, and 0.07–0.13, respectively, while reservoir size was fixed at 2,000 neurons. Forecast trajectories that produced non-finite values or exceeded a predefined instability threshold (10^6^ individuals), used solely to identify divergent simulations, were discarded, and predictive distributions were summarized from the remaining valid trajectories. When observed abundances are available for comparison, the workflow concludes with an independent assessment of forecast accuracy, predictive uncertainty, and forecast reliability, as described in the following sections.

The `reserBUGS` framework has been implemented as an open-source Python library to facilitate integration into existing ecological forecasting workflows [27]. The library supports construction of training datasets from population abundance records and sampling locations, and automatically retrieves environmental predictors through the official Copernicus and NASA Earthaccess APIs [38, 39]. Ensemble forecasts are generated using the reservoir computing implementation provided by Trouvain et al. [40], which is integrated into the framework to produce predictive distributions and associated confidence intervals.

### Dataset

We evaluated forecasting performance using empirical insect population time series derived from BIOTIME 2.0 [28]. The analysis was restricted to terrestrial species belonging to the class Insecta. Taxonomic identities were standardized using the GBIF Backbone Taxonomy [41]. Because sampling grain and spatial extent varied among BIOTIME studies, abundance records were harmonized prior to analysis. Observations from the same study were aggregated into standardized spatial units at a resolution of 1 ˣ 1 km, providing a consistent representation of local population dynamics across datasets.

We retained only time series that satisfied the following criteria: (i) species-level taxonomic identification, (ii) abundance data expressed as counts, (iii) consecutive annual observations, (iv) a minimum duration of ten years, and (v) at least one non-zero abundance observation. The final dataset comprised 2,858 species–location time series representing 1,018 insect species across nine studies and 36 sampling locations. Population abundance was modelled independently for each species–location time series. Time series were distributed among four major insect orders: Lepidoptera (75.0%), Hymenoptera (13.5%), Diptera (7.4%), and Coleoptera (4.1%). Study sites were concentrated in three broad geographic regions: East Asia, Europe, and North America (Supplementary Figure S1), encompassing a range of temperate, boreal, and semi-arid environments. This spatial and environmental heterogeneity provides a suitable test bed for evaluating the generality and robustness of ecological forecasting models across contrasting ecological contexts.

#### Environmental predictors

Environmental covariates were obtained from the Copernicus Climate Change Service (C3S) ERA5 reanalysis products. Predictor variables included total precipitation (m), air temperature (K), zonal and meridional wind components (m s*^−^*^1^), and annual mean leaf area index (LAI), a measure of vegetation canopy density (m^2^ leaf area per m^2^ ground area), for high (trees) and low (shrubs and herbs) vegetation. These variables represent major environmental drivers that can influence insect population dynamics through their effects on thermal conditions, moisture availability, vegetation structure, and dispersal processes [33, 42].

Monthly environmental data were aggregated to annual resolution to match the temporal resolution of the abundance records. Annual precipitation was calculated as the sum of monthly values, whereas all other environmental variables were averaged across months. Although species-specific phenological windows may capture stronger environmental signals for some taxa [43], we used annual aggregation to prioritize generality and comparability by providing a consistent predictor set across the diverse taxa and monitoring programs included in this study.

We additionally evaluated the potential value of the Normalized Difference Vegetation Index (NDVI) as a predictor of insect abundance using a random subset of 100 time series. Spearman correlations calculated at monthly and annual temporal resolutions (Supplementary Figure S2) revealed moderate to strong associations between NDVI and several environmental predictors already included in the forecasting models, particularly precipitation and vegetation-related variables. Given this overlap and the substantial computational cost of retrieving NDVI data for the full dataset, NDVI was not included in subsequent analyses.

#### Train and test sets

Time series were ordered chronologically. For each series, the final five years were reserved as an independent test period, while all preceding observations were used for model fitting. This rolling forecast horizon allowed model performance to be evaluated across prediction horizons ranging from one to five years ahead, thereby quantifying the propagation of forecast error and uncertainty through time.

#### Recursive forecasting and predictive distributions

Forecasts were generated recursively. Predictions for the first forecast year were produced using information available from the training period, and each predicted value was subsequently used as input for the next forecast step. This procedure was repeated until forecasts had been generated for the complete five-year test horizon. This recursive formulation implies that abundance depends jointly on previous abundance and environmental conditions.

To characterize predictive uncertainty, each model generated an ensemble of 500 forecast trajectories. These ensembles were used to approximate predictive distributions for abundance at each forecast horizon. We used the median of each predictive distribution as the point forecasts, a single representative prediction of future abundance, while the distribution as a whole characterized predictive uncertainty. This uncertainty reflects variation arising from the ensemble forecasting procedure, specifically through stochastic reservoir initialization and hyperparameter sampling within the adopted reservoir-computing framework. It should therefore be interpreted as conditional on the modelling framework and does not explicitly account for other important sources of uncertainty, including observation error, structural model uncertainty, or uncertainty associated with future environmental drivers [44].

These predictive distributions formed the basis for all subsequent analyses of forecast accuracy, probabilistic performance, and forecast reliability.

### Forecasting models

The forecasting framework was designed to evaluate the ability of competing modelling approaches to predict future abundance dynamics and quantify predictive uncertainty under a common evaluation protocol. To that end, we compared a hierarchy of forecasting approaches representing increasing levels of complexity and different assumptions about the processes governing population dynamics. This comparison framework allowed us to evaluate the relative importance of temporal autocorrelation, environmental information, nonlinear responses, and reservoir-based dynamic representations for ecological forecasting. Supplementary Table S1 summarizes the information available to each model, its general formulation, the approach used to generate probabilistic forecasts, and the proportion of time series yielding stable forecast ensembles.

#### Benchmark models

We first implemented three simple benchmark models that provide reference levels of forecasting performance and represent increasingly informative assumptions about future population dynamics.

The *mean model* assumes that abundance fluctuates around a constant long-term average according to

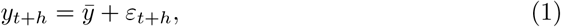

where *y_t_*_+*h*_ denotes the abundance predicted *h* time steps ahead from the last observation in the training period, *ȳ* is the mean abundance observed during the training period and *ε_t_*_+*h*_ is an independent random error term. This model ignores temporal structure and serves as a baseline expectation based solely on average historical conditions.

The *persistence model* assumes that future abundance remains centred on the most recent observation while allowing random fluctuations over time,

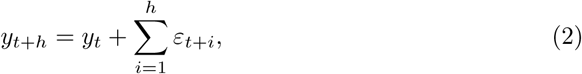

where *y_t_* denotes abundance at time *t*, taken here as the last observed value in the training period, *ε_t_*_+*i*_ represents the independent random error at time *t* + *i*, and their cumulative sum captures the cumulative stochastic variation over the forecast horizon. This model represents a widely used benchmark in ecological forecasting and provides a simple description of short-term population inertia.

The *random walk with drift model* (RW+drift) extends the persistence assumption by incorporating a constant mean annual change estimated from the training data,

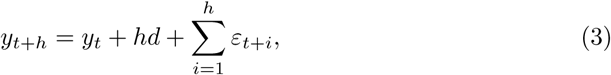

where *d* is the mean annual change in abundance during the training period; all other terms are as defined for the persistence model. This model therefore accounts for persistent directional changes while retaining a simple stochastic formulation.

Together, these benchmark models establish lower-bound expectations for forecasting performance against which more complex approaches can be evaluated.

#### Autoregressive models

To evaluate the predictive value of temporal dependence in abundance dynamics, we implemented two autoregressive forecasting approaches based exclusively on historical abundance observations [45].

The autoregressive model of order one (AR(1)) represents the simplest autoregressive description of population dynamics, in which abundance at time *t* depends linearly on abundance in the previous year:

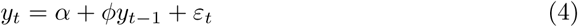

where *α* is the intercept, *ϕ* is the autoregressive coefficient, and *ε_t_* is a random error term. This model captures short-term temporal dependence while assuming linear population dynamics.

To allow for more complex temporal structures, we also fitted Auto-ARIMA models [45, 46], in which the orders of the autoregressive (*p*), differencing (*d*), and moving-average (*q*) components were selected automatically by minimizing the Akaike Information Criterion (AIC) (the mathematical formulation is provided in Appendix S1). As in AR(1), the autoregressive term relates future abundance to previous observations, whereas differencing captures changes through time rather than absolute abundance values, and the moving-average term accounts for temporal structure in past prediction errors. The automated search was restricted to non-seasonal models with autoregressive and moving-average orders up to three (*p* ≤ 3, *q* ≤ 3) and differencing orders up to two (*d* ≤ 2), using a stepwise search procedure to reduce computational cost.

Together, these models provide autoregressive benchmarks of increasing complexity while relying exclusively on historical abundance information, allowing the added value of reservoir-based forecasting to be evaluated against conventional temporal models.

#### Environmental forecasting models

To evaluate whether reservoir computing provides predictive advantages beyond those obtainable from conventional predictor-based models, we implemented two forecasting approaches based on the same environmental covariates and lagged abundance supplied to the reservoir models.

The *Tweedie generalized linear model* (GLM) combined environmental covariates with one-year lagged abundance using a log link and a Tweedie response distribution [37] (the mathematical formulation is provided in Appendix S2). This model captures additive relationships between predictors and abundance while assuming a linear predictor on the log scale.

We also implemented a *quantile random forest model* [47] using the same set of environmental predictors and lagged abundance (the mathematical formulation is provided in Appendix S3). Unlike the GLM, random forests can capture nonlinear responses and interactions among predictors without requiring a predefined functional form, providing a flexible machine-learning benchmark for evaluating the added value of reservoir-based dynamic representations. Predictive uncertainty was quantified directly from conditional quantiles estimated across the ensemble of decision trees, allowing construction of predictive distributions comparable to those generated by the reservoir models [47].

Comparisons between these models and the reservoir computing approach evaluate whether reservoir-based dynamic representations improve forecasting performance beyond conventional statistical and machine-learning approaches supplied with the same environmental predictors and lagged abundance information.

#### Reservoir computing models

The *reservoir computing* (RC) models used the same predictor set as the environmental forecasting models, comprising environmental covariates and one-year lagged abundance. A single lagged abundance predictor was used throughout the primary analyses to ensure comparable autoregressive information across forecasting approaches while maximizing the effective training sample size for the relatively short time series (at least 10 annual observations). The influence of additional lagged abundances was evaluated separately in the hyperparameter sensitivity analysis. Environmental covariates and lagged abundance jointly drove the reservoir dynamics, generating high-dimensional internal states that encoded recent system dynamics (the mathematical formulation is provided in Appendix S4). Expected abundance was then obtained by fitting a Tweedie generalized linear model with a log link to these reservoir states.

We implemented two reservoir variants. The *Full Reservoir* used both environmental covariates and one-year lagged abundance as dynamic inputs throughout the forecasting procedure, thereby integrating biotic memory and environmental variability. In contrast, the *Basic Reservoir* retained the same reservoir architecture but replaced the environmental covariates with a single constant dummy input (equal to one). Consequently, forecasts were driven exclusively by historical abundance dynamics and represent predictions based on internal population dynamics alone. This design isolated the contribution of environmental information while keeping the reservoir architecture unchanged.

### Forecast evaluation

We evaluated forecasting performance using both magnitude accuracy and probabilistic forecast performance. This combination allowed us to assess not only how accurately models predicted future abundance, but also how well the predictive distributions represented uncertainty in those predictions.

To ensure that differences in forecasting performance reflected differences among models rather than differences in the time series being evaluated, direct model comparisons were restricted to time series for which forecasts were available from all models.

#### Magnitude accuracy

Forecast accuracy was quantified using Type-M error, which measures multiplicative deviations between predicted and observed abundance on a logarithmic scale. For an individual prediction,

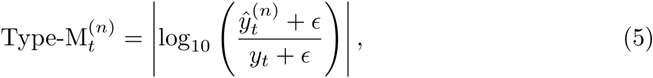

where *ŷ_t_^(n)^* is the abundance predicted by ensemble member *n*, *y_t_*is the observed abundance at time *t*, and *ɛ* = 1 is a small constant added to avoid instability when abundance values approach zero.

For each forecast horizon, Type-M error was calculated for each of the 500 forecast trajectories generated by a model. The resulting ensemble distribution of Type-M values was summarized using its median and interquartile range. Median Type-M values were used for model comparisons and skill score calculations.

#### Probabilistic forecast performance

Because all forecasting models were implemented to generate predictive distributions, we additionally evaluated forecast quality using proper scoring rules that assess the entire predictive distribution rather than only a point estimate [48].

We quantified probabilistic forecast performance using the Continuous Ranked Probability Score (CRPS), which measures the agreement between the predictive distribution and the observed outcome [48]. Lower CRPS values indicate more accurate and better calibrated probabilistic forecasts. Although the `reserBUGS` framework also supports the calculation of additional probabilistic scoring rules and uncertainty metrics, such as the Dawid–Sebastiani Score and interval score, CRPS was used as the primary probabilistic evaluation metric because it is a widely adopted scoring rule that provides an overall assessment of probabilistic forecast performance [48, 49].

#### Skill scores

To facilitate comparison across models and time series, we converted Type-M error and probabilistic scores into skill scores relative to a baseline forecasting model. Skill scores express forecasting performance as a proportional improvement relative to a common reference forecast, allowing results to be compared across heterogeneous time series and evaluation metrics [48, 50]:

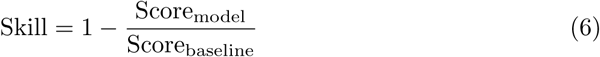

Positive skill values indicate that a model outperformed the baseline, whereas negative values indicate poorer performance.

#### Performance across forecast horizons

Forecast performance was evaluated separately for prediction horizons ranging from one to five years ahead. This allowed us to quantify how forecast accuracy and uncertainty changed as predictions extended further into the future.

Performance metrics were first calculated for each species–location time series and forecast horizon and subsequently summarized across all time series using medians and interquartile ranges.

### Forecast performance diagnostics

Beyond comparing average forecast performance across models, we asked whether characteristics of the predictive distribution could identify forecasting conditions under which the Full Reservoir was more likely to outperform or underperform a simpler autoregressive benchmark. To address this question, we characterized each forecast using two complementary diagnostics derived directly from the predictive distribution and therefore not requiring test observations: forecast displacement (*D*), which measures the extent to which forecasts depart from recent historical conditions, and uncertainty expansion (*U*), which quantifies whether predictive uncertainty exceeds the variability historically observed in the time series. Forecast displacement (*D*) and uncertainty expansion (*U*) captured the most informative properties of the predictive distribution when compared with a broader set of distributional descriptors, including measures of predictive spread, tail behaviour, asymmetry, entropy, multimodality, and distributional complexity. Statistical methods and complete results are provided in the Supplementary Material S5.

For each time series, we calculated forecast displacement as

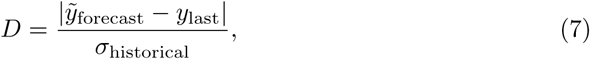

where *ỹ*_forecast_ is the median of the predictive distribution, *y*_last_ is the final observed abundance in the training period, and *σ*_historical_ is the standard deviation of historical abundances. Larger values indicate forecasts that project population states increasingly different from the most recently observed state relative to the historical variability of the time series.

Predictive uncertainty expansion was quantified as

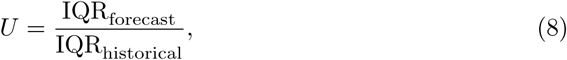

where IQR_forecast_ is the interquartile range of the predictive distribution and IQR_historical_ is the interquartile range of abundances observed during the training period. Values greater than one indicate that forecast uncertainty exceeds the variability historically observed in the population.

To identify forecasting conditions under which the Full Reservoir was more or less likely to outperform the AR(1) benchmark, we compared the predictive performance of both models at the first forecast horizon using the Continuous Ranked Probability Score (CRPS). Forecasts were assigned to two-dimensional bins defined by forecast displacement (*D*) and uncertainty expansion (*U*), using logarithmically spaced bins along both axes. Within each bin, we estimated the empirical probability of the Full Reservoir being outperformed by AR(1) as the proportion of time series for which its CRPS was at least 5% higher than that of AR(1). To provide an interpretable reference for these diagnostics, we additionally compared this probability between forecasts for which both *D* and *U* exceeded one and those for which both metrics were equal to or below one. To avoid unstable estimates arising from sparse data, bins containing fewer than eight forecasts were omitted. The resulting probabilities were visualized as relative performance maps, in which high values indicate forecasting conditions where AR(1) is more likely to outperform the Full Reservoir, whereas low values indicate conditions where the Full Reservoir is more likely to retain its predictive advantage.

### Sensitivity of reservoir forecasting performance to hyperparameter selection

To evaluate the sensitivity of the Full Reservoir model to hyperparameter selection, we performed an additional analysis in which multiple reservoir configurations were compared for selected time series. The hyperparameter grid explored combinations of reservoir size (100, 250, 500, and 2000 nodes), number of lagged observations (1, 2, and 3 years), spectral radius (0.7, 0.9, and 1.1), and leaking rate (0.3, 0.6, and 1.0), yielding 108 candidate configurations per time series. Collectively, these parameters determine the capacity of the reservoir to represent historical population dynamics and retain information through time [34].

Because evaluating all candidate configurations across the full dataset would have required fitting more than 300,000 reservoir models, the analysis was conducted on a random subset of 100 time series. For each series, the configuration with the lowest one-year-ahead CRPS was retained as the optimized Full Reservoir configuration. These optimized models should be interpreted as an oracle benchmark, as model selection was based on observed forecasting performance. The performance of the optimized and default Full Reservoir configurations was then compared using Type-M error and probabilistic skill metrics.

### Software and reproducibility

All analyses were conducted in Python 3.12.13 using the open-source scientific computing libraries NumPy 2.4.3, pandas 3.0.2, scikit-learn 1.8.0, statsmodels 0.14.6, pmdarima 2.1.1, and quantile-forest 1.4.1. The `reserBUGS` forecasting framework evaluated in this study is available as an open-source Python package [27] (https://github.com/JJ-Lab/reserBUGS). The study-specific code used to apply `reserBUGS` to the BioTIME dataset, implement the benchmark models, and reproduce all analyses and figures presented in this study is publicly available on GitHub (https://github.com/JJ-Lab/reserBUGS-biotime-forecast) and permanently archived on Zenodo (https://doi.org/10.5281/zenodo.21916497). The latter repository includes a portable Conda environment file (environment.yml) containing the direct dependencies required to reproduce the analyses, together with a complete snapshot of the software environment used in the study (environment full.yml).

ChatGPT (OpenAI) was used during manuscript preparation to assist with code development, debugging, and language refinement. All outputs were reviewed and validated by the authors, who are solely responsible for the ideas, analyses, interpretations, and conclusions presented in this manuscript.

## Results

The initial dataset comprised 2,858 insect abundance time series. To ensure that all forecasting approaches were evaluated on an identical set of observations, we restricted subsequent model comparisons to time series for which every model generated complete ensembles of 500 forecast trajectories across all five forecast horizons. This criterion was satisfied by 1,094 time series (Supplementary Table S1 and Supplementary Figures S3 and S4).

We found substantial differences among forecasting approaches in their ability to generate complete forecast ensembles. Mean, persistence, random walk with drift, and AR(1) generated complete ensembles for all time series, whereas Random Forest, Auto-ARIMA, Full Reservoir, Basic Reservoir, and Tweedie GLM achieved complete ensemble generation for 97.9%, 96.5%, 84.8%, 74.8%, and 55.4% of time series, respectively (Supplementary Table S1). Overlap analyses revealed that most exclusions from the common comparison dataset were attributable to the Tweedie GLM. In particular, an additional 822 time series generated complete ensembles for all models except Tweedie, identifying this model as the principal source of forecast failures (Supplementary Figures S3 and S4). These failures primarily resulted from occasional numerical overflows during recursive prediction, producing unrealistically large abundance estimates and preventing complete forecast ensembles from being generated.

We also observed differences between the two reservoir implementations. Compared with the Basic Reservoir, the Full Reservoir generated complete forecast ensembles for an additional 286 time series (2,424 versus 2,138), suggesting that inclusion of environmental predictors increased the proportion of series for which complete probabilistic forecasts could be obtained.

To assess whether this filtering altered the structure of the dataset, we compared the taxonomic and geographic composition of the common comparison subset with that of the full dataset. The retained subset comprised 1,094 time series (38.3% of the original dataset), representing 664 of the 1,018 species, seven of the nine studies, and 31 of the 36 sampling locations included in the full dataset. We found that the common comparison subset remained broadly representative of the original dataset (Supplementary Figures S5 and S6). Lepidoptera remained the dominant order, accounting for 76.5% of retained time series compared with 81.9% in the full dataset, whereas Hymenoptera increased from 11.1% to 14.1% and Diptera from 2.7% to 4.9%. Geographic representation was similarly preserved, with Europe, East Asia, and North America accounting for 51.0%, 35.1%, and 13.9% of retained time series, respectively. Together, these results indicate that restricting analyses to time series with complete forecast ensembles did not substantially alter the taxonomic or geographic structure of the dataset.

### Does reservoir computing improve ecological forecasting performance?

To evaluate whether reservoir computing improves ecological forecasting performance relative to established forecasting approaches, we first compared one-year-ahead forecasts using Type-M error and probabilistic skill metrics (Figure 2). Skill scores were calculated relative to the mean model because persistence-based skill scores were strongly distorted in a subset of time series where the final training observation was close to zero, producing unstable denominators and extreme skill values.

**Fig 2.**
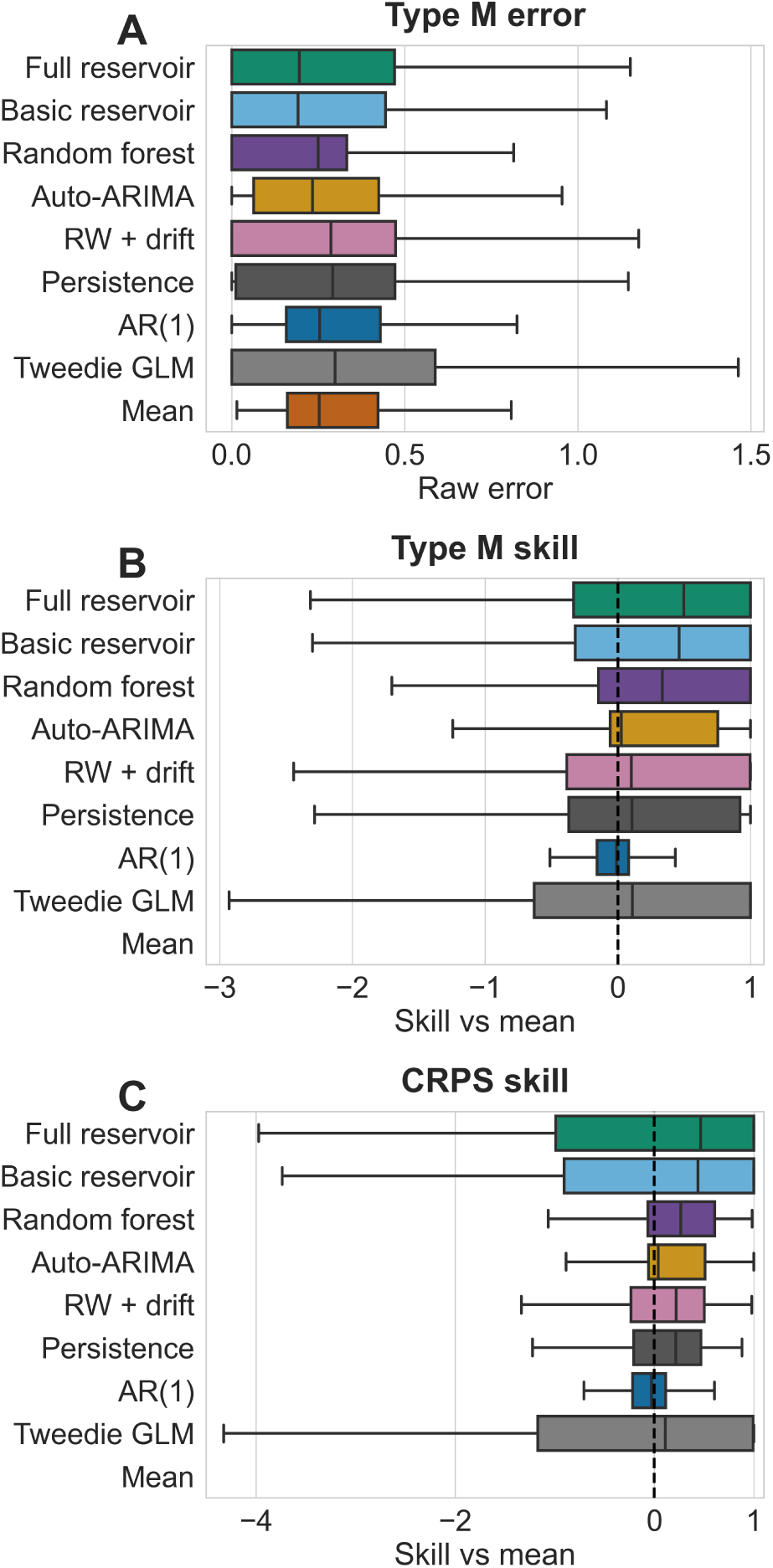
One-year-ahead forecasting performance across models. A) Distribution of Type-M error, B) Type-M skill, and C) CRPS skill for one-year-ahead forecasts. Type-M error quantifies the discrepancy between predicted and observed abundance on a logarithmic magnitude scale, with lower values indicating better performance. Skill scores are calculated relative to the baseline model; positive values indicate improvement over the baseline. Boxes show the interquartile range, central lines indicate medians, and whiskers show the range of values across time series.

Across models, both reservoir approaches consistently ranked among the best-performing methods. The Basic Reservoir achieved the lowest median Type-M error at the one-year-ahead horizon (0.18), followed closely by the Full Reservoir (0.20), indicating that predicted changes in abundance typically differed from the observed magnitude of population change by 18–20%. Random forest also performed well, although with slightly larger median errors than the reservoir approaches. In contrast, benchmark and autoregressive models typically exhibited median errors between 0.25 and 0.30 (Figure 2A). The Tweedie GLM showed substantially greater variability among time series, with the widest interquartile range across all three performance metrics despite median performance comparable to several benchmark models.

When performance was evaluated relative to the mean model, reservoir approaches again achieved the highest skill scores. The Full Reservoir produced the highest median Type-M skill (0.50), followed by the Basic Reservoir (0.47), indicating approximately a 50% reduction in magnitude error relative to the baseline forecast (Figure 2B). Random Forest ranked third (0.34), whereas autoregressive and benchmark models provided little or no improvement over the baseline. Unlike most other models, both reservoir models combined high median skill with relatively compact interquartile ranges, indicating that their performance advantages were consistently observed across a large proportion of time series.

A similar pattern emerged for probabilistic performance. The Full Reservoir achieved the highest median CRPS skill (0.46), followed by the Basic Reservoir (0.26) and Random Forest (0.25) (Figure 2C). These positive skill scores indicate that reservoir-based forecasts produced more accurate and better calibrated predictive distributions than the mean baseline forecast.

### How does forecasting performance change with prediction horizon?

Forecast performance showed little evidence of systematic deterioration with increasing prediction horizon. Reservoir-based approaches consistently outperformed benchmark, autoregressive, and environmental forecasting models across the full five-year forecasting horizon (Figure 3). Notably, both reservoir models exhibited a slight improvement in Type-M error and probabilistic skill at intermediate horizons (3–4 years). The Full Reservoir generally outperformed the Basic Reservoir, with the largest performance differences occurring at these intermediate horizons, suggesting that environmental predictors contributed additional predictive information beyond historical abundance dynamics alone.

**Fig 3.**
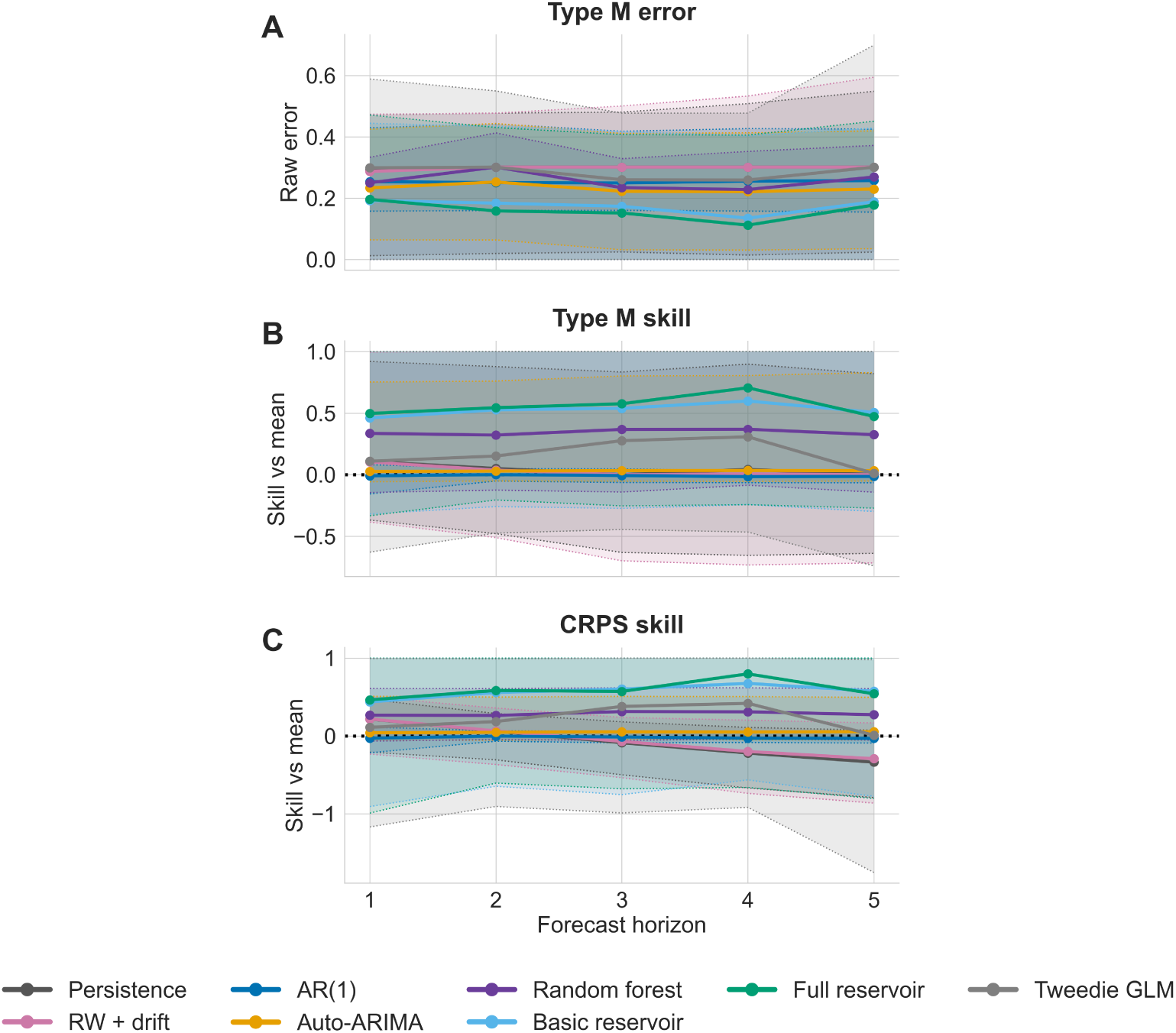
Forecasting performance across the five-year prediction horizon. A) Median Type-M error, B) Type-M skill, and C) CRPS skill are shown for each model from one- to five-year-ahead forecasts. Shaded areas represent uncertainty around the across-series summary. Lower Type-M error indicates more accurate magnitude forecasts, whereas higher skill values indicate stronger improvement relative to the baseline model.

### Can probabilistic forecasts identify conditions associated with reduced predictive skill?

Although the Full Reservoir achieved the best median forecasting performance across metrics and forecast horizons (Figures 2 and 3), its predictive skill varied substantially among individual time series. Forecasts characterized by larger departures from recent historical conditions (*D*) and greater uncertainty expansion (*U*) showed reduced performance relative to AR(1). Both diagnostics were positively correlated with the relative CRPS of the Full Reservoir compared with AR(1) (*D*: Spearman’s *ρ* = 0.55; *U* : *ρ* = 0.59; both *P <* 10*^−^*^180^).

This pattern was particularly pronounced when both diagnostics exceeded one. Forecasts with *D >* 1 and *U >* 1 had an 88% probability of the Full Reservoir having a CRPS at least 5% higher than AR(1), compared with 32% when both metrics remained at or below one. When only one diagnostic exceeded one, this probability was intermediate (51% for *D >* 1 and *U* ≤ 1, and 64% for *D* ≤ 1 and *U >* 1).

Consistently, the probability of the Full Reservoir being outperformed by AR(1) increased as forecasts moved further away from the recent historical state (*D*) and as predictive uncertainty (*U*) expanded relative to the historical variability of the time series (Figure 4). Regions characterized by small forecast displacement and uncertainty comparable to historical variability were associated with lower probabilities of AR(1) outperforming the Full Reservoir, whereas large displacement combined with expanded predictive uncertainty was associated with substantially higher probabilities. These results show that properties of the predictive distribution itself can provide diagnostic information about the conditions under which the predictive advantage of the Full Reservoir is more likely to deteriorate.

**Fig 4.**
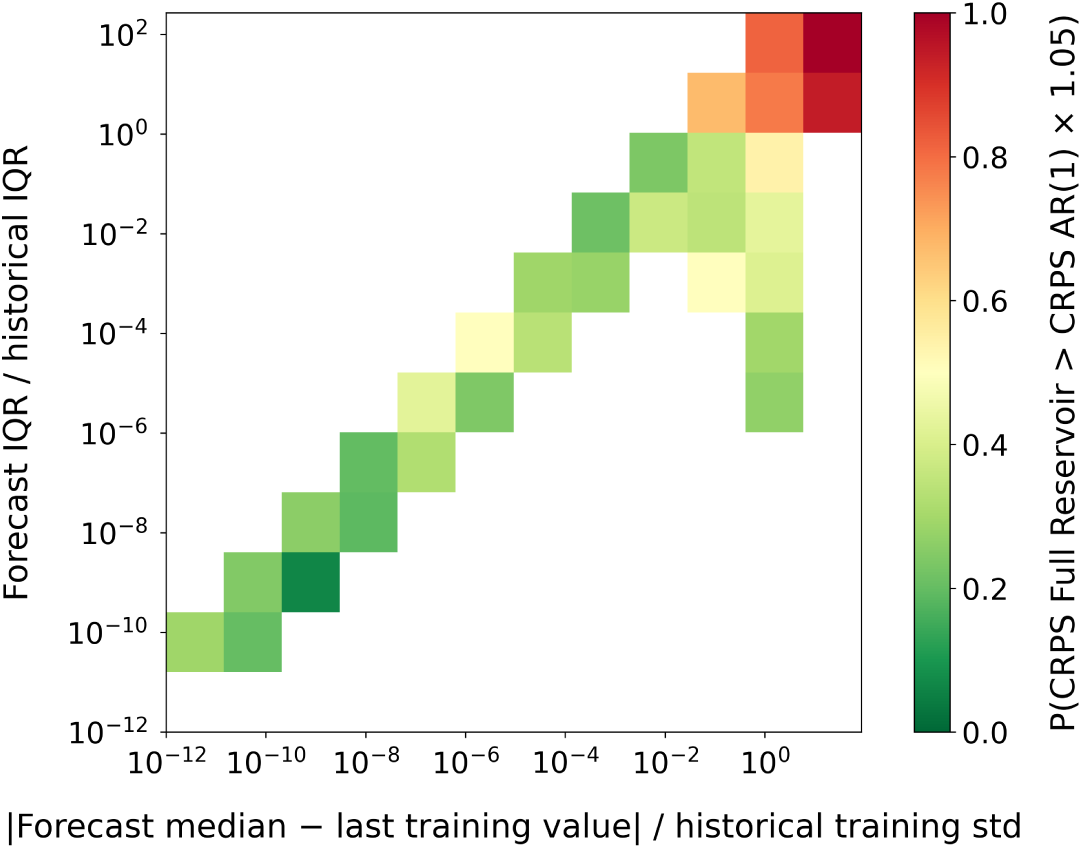
Forecast performance map for the Full Reservoir model. Forecasts were characterized by two properties of the predictive distribution: the standardized displacement of the forecast median from the last observed training value (*D*; x-axis) and the relative predictive uncertainty (*U* ; y-axis), measured as the ratio between forecast and historical interquartile ranges. Colours indicate the empirical probability that the Full Reservoir produced a CRPS at least 5% worse than the AR(1) model. White cells indicate bins containing fewer than eight forecasts, for which empirical probabilities were not estimated. Green cells correspond to regions of forecast space where the reservoir was rarely outperformed by AR(1), whereas red cells indicate conditions associated with a high risk of poorer performance. Relative forecast performance declined with increasing forecast displacement (*D*; Spearman’s *ρ* = 0.55, p-value *<* 10*^−^*^180^) and uncertainty expansion (*U* ; *ρ* = 0.59, p-value *<* 10*^−^*^180^). Forecasts with both *D >* 1 and *U >* 1 had an 88% probability of the Full Reservoir producing a CRPS at least 5% higher than AR(1), compared with 32% when both metrics remained at or below one.

### How sensitive is reservoir forecasting performance to hyperparameter selection?

To assess the sensitivity of reservoir forecasting performance to hyperparameter selection, we compared the original reservoir configuration with a series-specific optimized reservoir. For each of 100 randomly selected time series, the optimized reservoir was defined as the configuration achieving the lowest one-year-ahead CRPS among the 108 reservoir configurations explored in the hyperparameter grid search.

The optimized reservoir consistently outperformed the original reservoir across all evaluated metrics (Figure 5). Median Type-M error decreased from approximately 0.30 to 0.11, indicating improved agreement between predicted and observed magnitudes of population change. Improvements were highly consistent across time series, with Type-M error decreasing in 98% of series and both Type-M and CRPS skill improving in 98–99% of cases.

**Fig 5.**
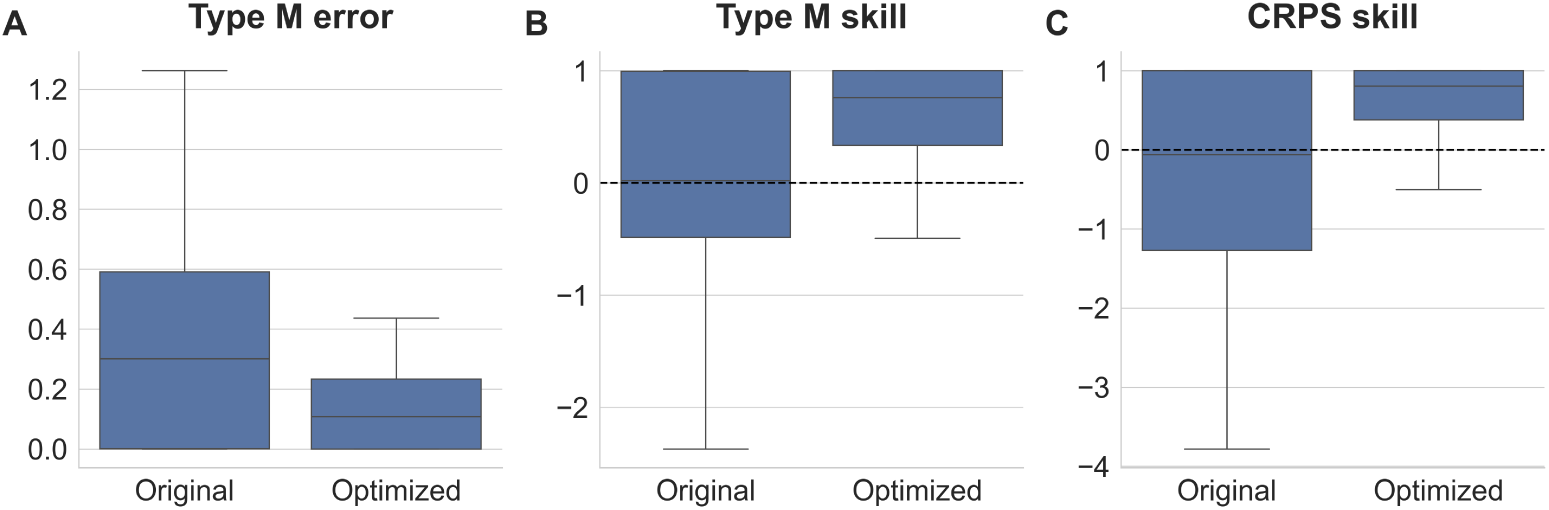
Sensitivity of reservoir forecasting performance to hyperparameter selection. Comparison between the original reservoir configuration and a series-specific optimized reservoir selected from a grid of 108 candidate hyperparameter combinations. Performance is shown for one-year-ahead forecasts using raw Type-M error (A), Type-M skill relative to the mean model (B), and CRPS skill relative to the mean model (C). Lower Type-M error indicates more accurate predictions of the magnitude of population change, whereas positive skill values indicate improvement relative to the baseline forecast. The optimized reservoir was selected independently for each time series using the configuration with the lowest one-year-ahead CRPS and therefore represents an oracle estimate of the maximum performance gain achievable through hyperparameter tuning.

Hyperparameter optimization also reduced variability among time series. Whereas the original reservoir exhibited a broad distribution of skill values, including many forecasts performing worse than the mean baseline, the optimized reservoir produced more compact distributions and predominantly positive skill values for both Type-M and CRPS metrics (Figure 5).

No single reservoir configuration emerged as universally optimal across the 100 time series included in the hyperparameter analysis. The most frequently selected configuration was optimal for only 9% of series, indicating substantial heterogeneity in optimal parameter combinations among forecasting problems. However, we can identify some patterns for individual hyperparameters. Larger reservoirs (2000 nodes) were selected most frequently, accounting for 56% of optimal configurations, whereas smaller reservoirs (250 or 500 nodes) were rarely selected. Similarly, spectral radii of 0.7 and 1.1 together accounted for 90% of the selected configurations, suggesting a preference for these values across forecasting problems. In contrast, the optimal number of lags and leaking rate showed no clear preference, with all tested values selected at comparable frequencies.

Because the optimal configuration was selected separately for each time series using observed forecasting performance, these results should be interpreted as an upper-bound benchmark representing the maximum performance achievable within the explored hyperparameter space rather than as an unbiased estimate of predictive skill.

## Discussion

Ecological forecasting faces a fundamental tension between model flexibility and data availability. Population abundance time series are typically short, noisy, and non-stationary [10], yet many modern forecasting approaches require large volumes of training data or extensive tuning to reach their full predictive potential. Here, we introduced `reserBUGS`, a reservoir-computing-based framework designed to navigate this tension. As an example, we have evaluated its performance on 2, 858 real insect abundance time series spanning multiple taxa, regions, and environmental contexts.

Our results demonstrate that reservoir-based approaches consistently outperformed established statistical and machine-learning benchmarks across both magnitude accuracy and probabilistic performance metrics. At the one-year-ahead horizon, the Full Reservoir reduced magnitude error by approximately 50% relative to the mean baseline, and this advantage was maintained across the full five-year forecasting window. Crucially, this improvement was not limited to point predictions: reservoir models also produced better-calibrated predictive distributions, as reflected in superior CRPS scores relative to autoregressive, GLM, and random forest alternatives. These findings suggest that the ability of reservoir computing to encode non-linear temporal dynamics and history-dependent effects within a computationally efficient architecture translates into meaningful forecasting gains under the data constraints typical of biodiversity monitoring.

Although the incorporation of environmental predictors further improved forecasting performance, particularly at intermediate forecast horizons of three to four years, the Basic Reservoir alone already constitutes a powerful ecological forecasting model, achieving error rates below 20% while consistently outperforming classic autoregressive approaches. Because it relies exclusively on historical abundance dynamics, it avoids the need for future values of environmental covariates, which must themselves be forecasted or assumed in ex-ante prediction [51], making it a practical and versatile option for broad implementation. Nevertheless, the Full Reservoir provided additional predictive skill beyond that contained in historical abundance dynamics alone. This result is consistent with the well-established influence of thermal conditions, moisture availability and vegetation structure on insect population dynamics [52], and highlights the potential of integrating remotely sensed and reanalysis-based environmental data into operational ecological forecasting workflows [8, 53]. Furthermore, the incorporation of environmental covariates improved forecast stability, with the Full Reservoir successfully generating complete forecast ensembles for substantially more time series than the Basic Reservoir, suggesting that environmental information not only enhances predictive accuracy but also reduces the risk of dynamical instability during recursive forecasting.

The improved performance of the Full Reservoir also revealed an important practical consideration: reservoir forecasts were sensitive to hyperparameter selection. Although this increases the complexity of model specification, it also suggests that the default configuration can be further improved through systematic optimization. Series-specific optimization reduced Type-M error from approximately 0.30 to 0.11 and improved skill scores in 98 − 99% of time series, demonstrating that substantial performance gains are achievable through tuning. However, no single configuration was universally optimal, and the most frequently selected configuration was best for only 9% of series. This heterogeneity underscores the need for principled hyperparameter selection strategies that go beyond fixed defaults. Adaptive or data-driven approaches to hyperparameter tuning, such as cross-validated grid search or Bayesian optimization, could help close the gap between the default and oracle configurations without relying on held-out performance as a selection criterion.

While these results are encouraging, several limitations warrant consideration. First, although `reserBUGS` was evaluated on a large and difficult taxonomically diverse dataset, the analysis was restricted to insect abundance time series derived from structured monitoring programs with annual resolution. Whether reservoir computing retains its advantages for taxa with slower life cycles or for time series at finer temporal resolutions (e.g. monthly sampling regimes) remains an open question. In principle, the reservoir computing framework is broadly applicable to any ecological group for which abundance or occurrence time series are available [28]. Reservoir dynamics can capture temporal dependencies across multiple timescales, including short-term fluctuations, longer-term memory effects, and potentially seasonal structure, through the leaking rate parameter, which controls the integration timescale of the reservoir memory [54]. This flexibility suggests that the framework could be extended to vertebrates, plants, or marine organisms, provided that sufficient time series are available to estimate the readout layer reliably. Systematic evaluation across a broader range of taxa and monitoring schemes would be a valuable direction for future work.

Second, the predictive advantage of the Full Reservoir was not uniform across the forecasting conditions. Although reservoir computing outperformed alternative approaches on average, some time series produced numerically unstable forecast trajectories, occasionally preventing complete forecast ensembles from being generated. This few, but large mispredictions might compromise its utility for real world conservation forecasts with conservation or policy implications [55]. Luckily, we provide simple metrics to identify such cases, and hence guide the users on assessing model performance. Specifically, we show that the probability that the Full Reservoir was outperformed by a simple AR(1) model increased markedly when forecasts projected large departures from recent historical conditions and when predictive uncertainty expanded relative to the historical variability of the time series. The performance map provides a practical tool for identifying forecasts that should be interpreted with additional caution, enabling practitioners to flag potentially unreliable predictions before they are used to inform management actions.

Third, although `reserBUGS` generates probabilistic forecasts, the predictive distributions should not be interpreted as a complete characterization of ecological uncertainty. Instead, they represent uncertainty arising from the forecasting procedure under the adopted reservoir-computing framework. Other important sources of uncertainty, including structural model uncertainty, observation error, and uncertainty in future environmental drivers, are not explicitly represented [44]. Consequently, our reservoir computing framework can be complemented by incorporating additional sources of uncertainty alongside the ensemble-based uncertainty considered here [3, 44].

A particularly promising direction for future development concerns the integration of `reserBUGS` into near-term iterative forecasting cycles. Near-term ecological forecasting, in which models generate short-horizon predictions that are evaluated against subsequent observations as they become available, has emerged as a powerful paradigm for adaptive management and cumulative learning across ecological systems [3, 25, 56]. The computational efficiency of reservoir computing, combined with the automated data retrieval and ensemble forecasting capabilities of the `reserBUGS` pipeline, makes the framework well-suited for deployment in operational monitoring contexts. As biodiversity monitoring programmes increasingly move towards higher-frequency data collection and open data sharing (e.g. [57]), the ability to generate probabilistic forecasts rapidly and at scale becomes a central requirement [3, 56]. Extending `reserBUGS` to support rolling forecast updates and automated performance tracking would represent a natural next step towards genuinely operational ecological forecasting.

In summary, `reserBUGS` provides a scalable, probabilistic, and computationally efficient framework for ecological forecasting that compares favourably with established alternatives across a diverse set of insect time series. Its integration of environmental predictors, ensemble uncertainty quantification, and forecast reliability assessment within a single open-source workflow addresses several practical barriers to the adoption of modern forecasting methods in biodiversity monitoring. We anticipate that reservoir computing, as implemented here, will prove useful not only for insect populations but more broadly as a flexible and data-efficient approach to forecasting ecological dynamics across taxa and systems.

