## Supporting information for "reserBUGS: A reservoir computing framework for probabilistic forecasting of ecological abundance time series"

#### Contents


- Table S1. Summary of forecasting models evaluated in the study.
- Figure S1. Geographic distribution of the 36 study locations included in the analysis.
- Figure S2. Spearman correlations among annual abiotic predictors and NDVI.
- Figure S3. Overlap among time series with complete forecast ensembles across forecasting models.
- Figure S4. Pairwise overlap in time series with complete forecast ensembles across forecasting models.
- Figure S5. Taxonomic composition of the full dataset and common comparison subset.
- Figure S6. Geographic composition of the full dataset and common comparison subset.
- Appendix S1. Autoregressive models
- Appendix S2. The Tweedie generalized model.
- Appendix S3. The quantile random forest model.
- Appendix S4. Reservoir computing models.
- Appendix S5. Forecast reliability analysis.
  - Table S2. Predictive distribution characteristics associated with relative forecast skill.
  - Table S3. Historical time-series characteristics associated with relative forecast skill.

**Table S1. Summary of forecasting models evaluated in the study.** The table reports the information used by each model, its general forecasting formulation, and the number of time series for which all 500 forecast trajectories remained valid across the five-year prediction horizon. Detailed mathematical descriptions and implementation details for the ARIMA, Tweedie GLM, Quantile Random Forest, and Reservoir Computing models are provided in Appendices S1–S4, respectively.

| **Model** | **Information used** | **Model form** | **Predictive distribution** | **Valid series** |
|---|---|---|---|---|
| Mean | Historical abundance mean | $y_{t+h} = \bar{y} + \varepsilon_{t+h}$ | Ensemble trajectories | 2858 (100.0%) |
| Persistence | Last observed abundance | $y_{t+h} = y_t + \sum_{i=1}^{h} \varepsilon_{t+i}$ | Ensemble trajectories | 2858 (100.0%) |
| RW + drift | Last abundance and historical trend | $y_{t+h} = y_t + hd + \sum_{i=1}^{h} \varepsilon_{t+i}$ | Ensemble trajectories | 2858 (100.0%) |
| AR(1) | Lagged abundance | $y_t = \alpha + \phi y_{t-1} + \varepsilon_t$ | Parametric forecast distribution | 2858 (100.0%) |
| Auto-ARIMA | Historical abundance series | ARIMA$(p, d, q)$ selected by AIC | Parametric forecast distribution | 2758 (96.5%) |
| Tweedie GLM | Lagged abundance and abiotic variables | Tweedie GLM with log link | Simulated forecast trajectories | 1583 (55.4%) |
| Random forest | Lagged abundance and abiotic variables | Quantile random forest | Conditional quantiles | 2798 (97.9%) |
| Basic Reservoir | Lagged abundance | Reservoir states with Tweedie readout | Ensemble trajectories | 2138 (74.8%) |
| Full Reservoir | Lagged abundance and abiotic variables | Reservoir states with Tweedie readout | Ensemble trajectories | 2424 (84.8%) |

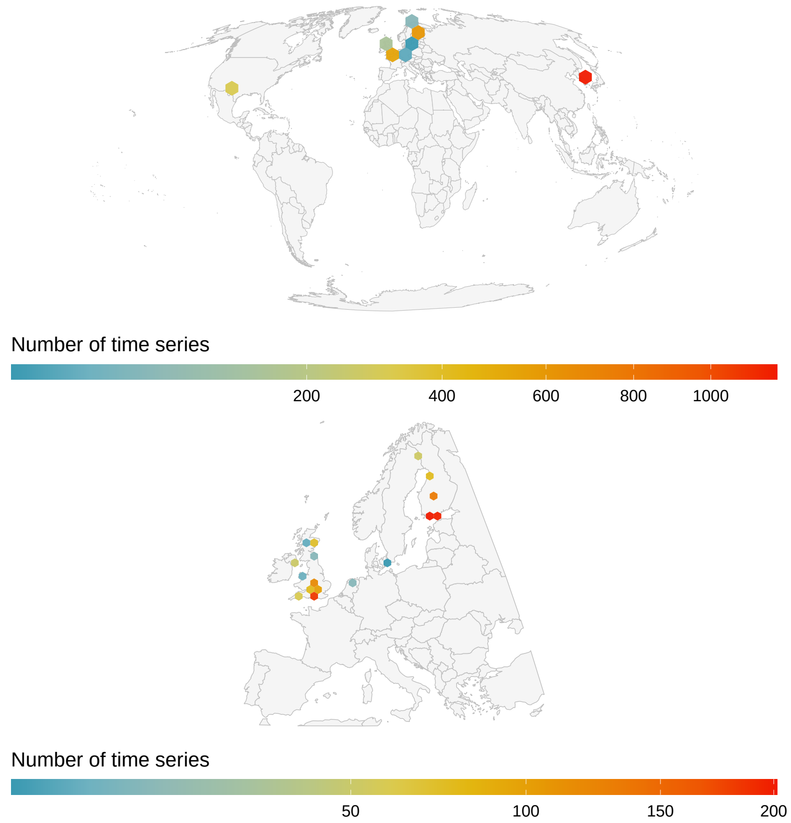


**Fig S1. Geographic distribution of the 36 study locations included in the analysis.** Locations are clustered across three main regions: East Asia (primarily South Korea), Europe (including the United Kingdom, Ireland, Central Europe, and Scandinavia), and North America (southwestern United States). Coordinates represent standardized sampling locations following spatial aggregation of population time series. Together, these sites span a range of temperate, boreal, and arid ecosystems.

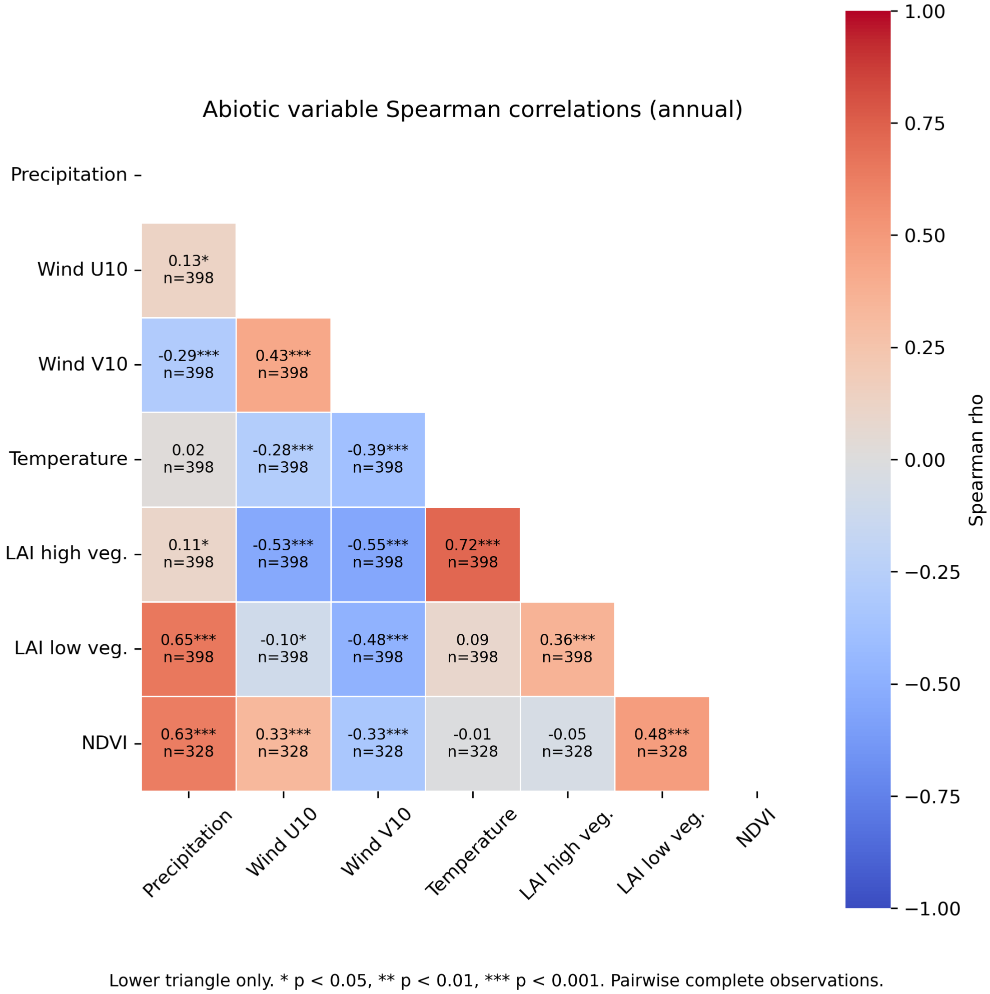


**Fig S2. Spearman correlations among annual abiotic predictors and NDVI.** Cells show pairwise Spearman rank correlation coefficients calculated from annual averages of environmental variables across the subset of time series used to evaluate the potential contribution of NDVI as an additional predictor. Values within cells indicate the correlation coefficient ($\rho$), sample size, and significance level. NDVI exhibited moderate to strong positive correlations with precipitation ($\rho = 0.63$) and low vegetation leaf area index (LAI; $\rho = 0.48$), indicating substantial overlap with environmental information already represented by existing predictors. Strong correlations were also observed among several vegetation and climate variables, particularly between temperature and high vegetation LAI ($\rho = 0.72$). Given the substantial redundancy between NDVI and predictors already included in the forecasting framework, together with the computational cost of obtaining NDVI data for all time series, NDVI was not included in subsequent forecasting analyses.

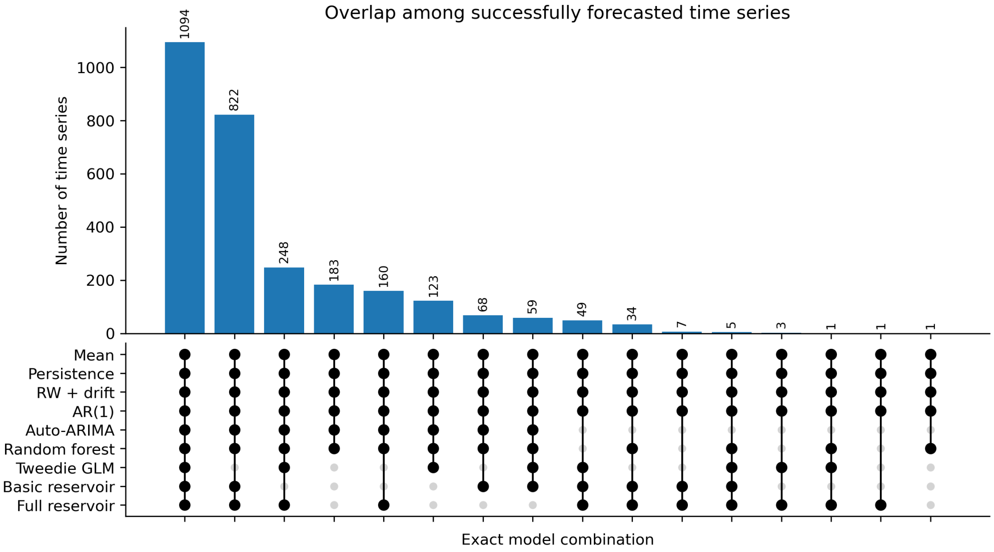


**Fig S3. Overlap among time series with complete forecast ensembles across forecasting models.** Bars represent exact intersections of time series for which forecasting models generated complete ensembles of 500 forecast trajectories across all five forecast horizons. The largest intersection comprised 1,094 time series for which all forecasting approaches produced complete ensembles. An additional 822 time series generated complete ensembles for all models except the Tweedie GLM, indicating that this model accounted for most reductions in the common comparison dataset. In contrast, the Full Reservoir generated complete ensembles for substantially more time series than the Basic Reservoir, highlighting differences in forecast stability among forecasting approaches.

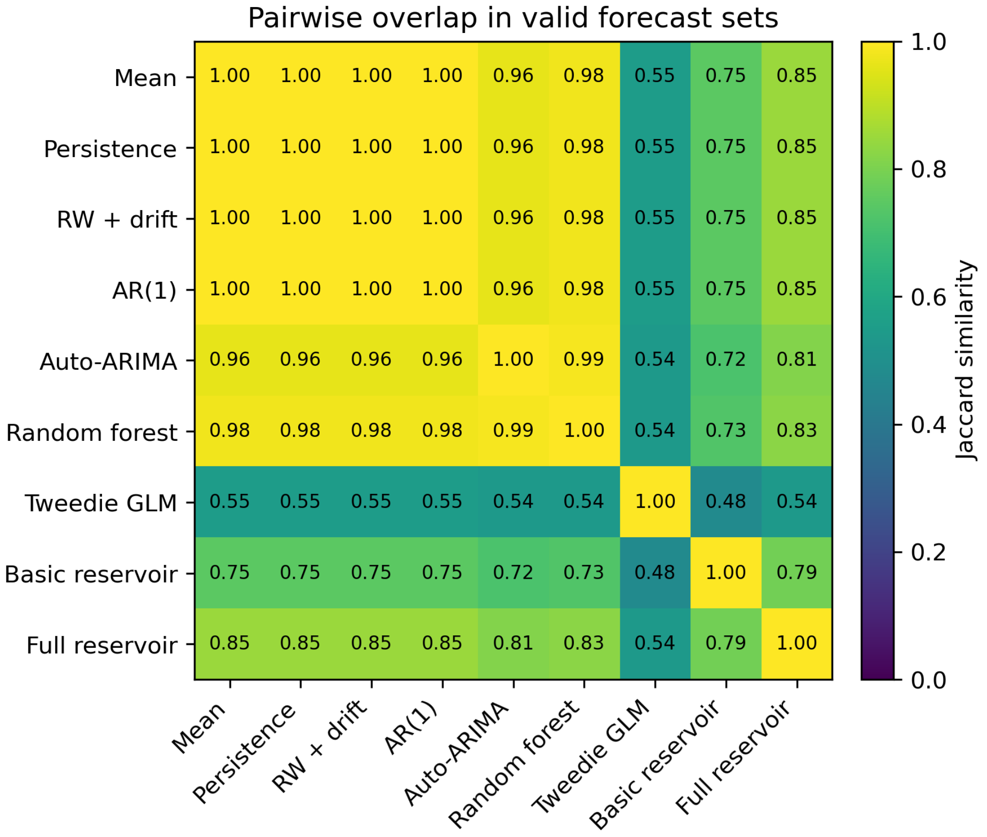


**Fig S4. Pairwise overlap in time series with complete forecast ensembles across forecasting models.** Cells show Jaccard similarity coefficients calculated from the sets of time series for which each model generated complete ensembles of 500 forecast trajectories across all five forecast horizons. Values close to 1 indicate that two models produced complete ensembles for largely the same set of time series, whereas lower values indicate greater differences in forecast coverage. Benchmark, autoregressive, and random forest models showed complete overlap (Jaccard = 1.0), while the lowest similarities involved the Tweedie GLM. The Full Reservoir exhibited greater overlap with the benchmark models than the Basic Reservoir, reflecting its ability to generate complete forecast ensembles for a larger number of time series.

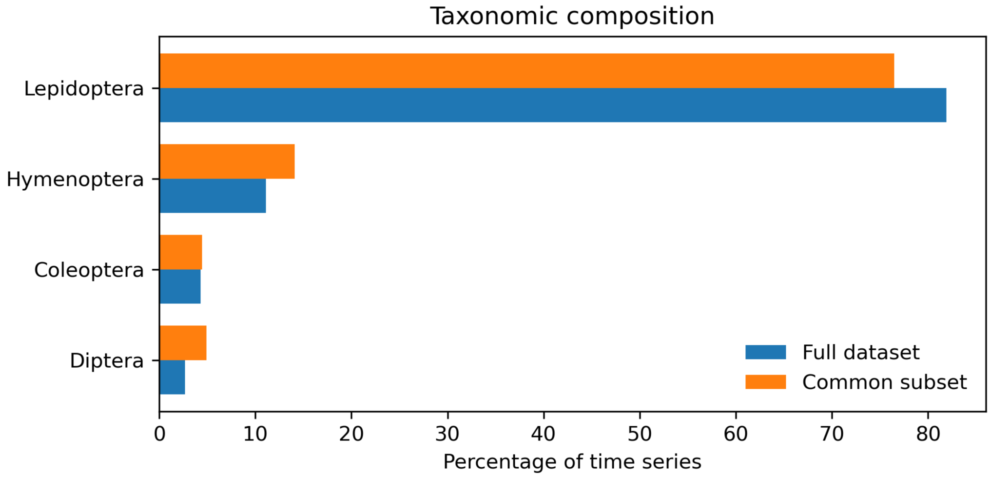


**Fig S5. Taxonomic composition of the full dataset and common comparison subset.** Bars show the percentage of time series belonging to each insect order in the full dataset (2,858 time series) and in the common comparison subset (1,094 time series) used for direct model comparisons. Lepidoptera remained the dominant order in both datasets, accounting for 81.9% of time series in the full dataset and 76.5% in the common comparison subset. Hymenoptera and Diptera were slightly overrepresented in the retained subset, whereas Coleoptera showed little change. Overall, restricting analyses to time series with complete forecast ensembles preserved the broad taxonomic structure of the original dataset.

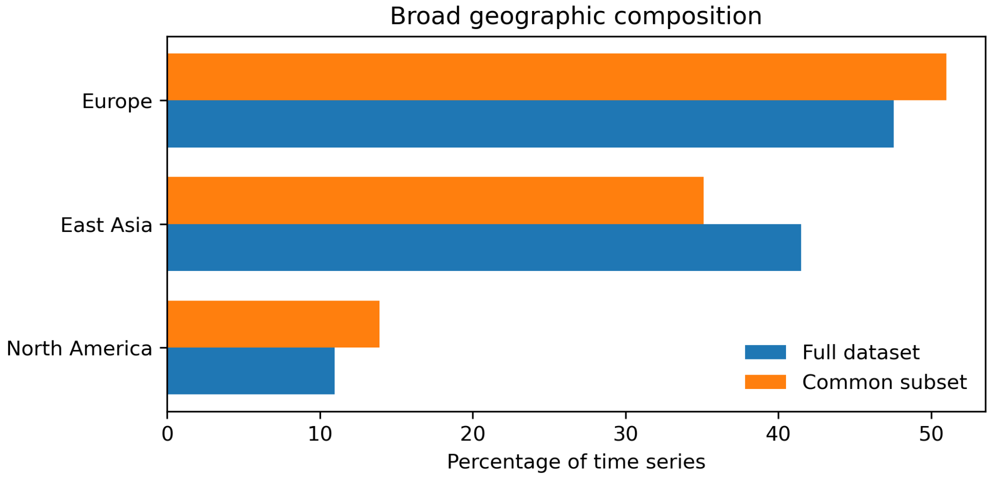


**Fig S6. Geographic composition of the full dataset and common comparison subset.** Bars show the percentage of time series originating from the three major geographic regions represented in the study: Europe, East Asia, and North America. Although the common comparison subset retained only time series for which all forecasting models generated complete forecast ensembles, regional representation remained broadly similar to that of the full dataset. Europe remained the most represented region, while East Asia was slightly underrepresented and North America slightly overrepresented in the retained subset. These results indicate that restricting analyses to complete forecast ensembles did not substantially alter the large-scale geographic composition of the dataset.

### S1 Autoregressive models.

While the autoregressive model of order one, AR(1), represents the simplest autoregressive description of population dynamics, in which abundance at time $t$ depends linearly on abundance in the previous year, models with a more complex temporal structures are also validated.

While the autoregressive term relates future abundance to previous observations, a differencing term captures changes through time rather than absolute abundance values and a moving-average term accounts for temporal structure in past prediction errors. The automated search was restricted to non-seasonal models with autoregressive and moving-average orders up to three ($p \leq 3$, $q \leq 3$) and differencing orders up to two ($d \leq 2$), using a stepwise search procedure to reduce computational cost.

$$\left(1 - \sum_{i=1}^{p} \phi_i B^i\right)(1-B)^d y_t = \alpha + \left(1 + \sum_{j=1}^{q} \theta_j B^j\right)\varepsilon_t, \tag{S1}$$

where $B$ is the backshift operator ($By_t = y_{t-1}$).

Together, these models provide autoregressive benchmarks of increasing complexity against which the forecasting performance of reservoir-based approaches can be evaluated, while relying exclusively on historical abundance information.

We also fitted Auto-ARIMA models [1, 2], in which the orders of the autoregressive ($p$), differencing ($d$), and moving-average ($q$) components were selected automatically by minimizing the Akaike Information Criterion (AIC).

### S2 The Tweedie generalized model.

The Tweedie generalized linear model (GLM) combined environmental covariates with one-year lagged abundance using a log link and a Tweedie response distribution [3]. The Tweedie distribution is well suited to ecological abundance data because it accommodates non-negative, right-skewed responses and can represent both exact zeros and continuous positive values within a single modelling framework. The expected abundance was modelled as:

$$\log(\hat{y}_t) = \beta_0 + \beta_{\text{lag}} \cdot y_{t-1} + \sum_{i=1}^{k} \beta_i X_{i,t}, \tag{S2}$$

where $\hat{y}_t$ is the expected abundance at time $t$ ($\hat{y}_t = \mathbb{E}\left[y_t | y_{t-1}, \mathbf{X}_t\right]$), $y_{t-1}$ is abundance in the previous year, and $X_{i,t}$ denotes the environmental predictors. This model captures additive relationships between predictors and abundance while assuming a linear predictor on the log scale.

We implemented the Tweedie GLM using a variance power parameter of 1.5 and a logarithmic link function. We applied ridge regularization ($\alpha = 10^{-4}$) and allowed a maximum of 10,000 optimization iterations. Continuous predictors were rescaled to the $[0, 1]$ interval before model fitting to improve numerical stability. To obtain probabilistic forecasts, we recursively simulated 500 abundance trajectories by sampling from the Tweedie distribution at each forecast step using the fitted conditional mean and the dispersion parameter estimated from the Pearson residuals. Predictive intervals and probabilistic forecast metrics were then computed from the resulting ensemble.

### S3 The quantile random forest model.

Like the Tweedie generalized linear model, the quantile random forest model used the environmental covariates together with the previous year's abundance as predictors,

$$\mathbf{x}_t = (X_{1,t}, X_{2,t}, \ldots, X_{k,t}, y_{t-1}), \tag{S3}$$

where $X_{i,t}$ denotes the environmental predictors and $y_{t-1}$ is abundance in the previous year.

Unlike the generalized linear model, the relationship between predictors and abundance was estimated non-parametrically using an ensemble of regression trees [4]. Each tree partitions the predictor space into terminal regions, and predictions are obtained from the observed training responses falling within the same terminal node as the new observation. Aggregating predictions across the ensemble yields an estimate of the conditional distribution

$$P(y_t \mid \mathbf{x}_t), \tag{S4}$$

rather than only its conditional mean.

Probabilistic multi-step forecasts were generated recursively. At each forecast horizon, one conditional quantile was sampled from the estimated distribution and the resulting abundance was used as the lagged predictor for the subsequent prediction step. Repeating this procedure generated an ensemble of forecast trajectories from which predictive intervals and other distributional summaries were obtained.

To implement the quantile random forest we used 500 decision trees with a minimum terminal node size of five observations and the default feature subsampling strategy of $\sqrt{p}$ predictors at each split, where $p$ denotes the total number of predictor variables. To generate probabilistic multi-step forecasts, at each forecast horizon $h$ we sampled a quantile level $q_h \sim U(0.01, 0.99)$ and evaluated the fitted conditional quantile function,

$$\hat{y}_h = \hat{Q}(q_h \mid \mathbf{x}_h).$$

The sampled abundance was recursively fed back as the lagged predictor for the next forecast horizon. We repeated this procedure to generate an ensemble of 500 forecast trajectories for each time series, from which predictive means and the 5th, 50th and 95th percentiles were computed.

### S4 Reservoir computing models.

Like the Tweedie generalized model, the reservoir computing model uses the abiotic covariates and the lagged abundance as the input vector $\mathbf{u}$ at time $t$, defined as

$$\mathbf{u}_t = (X_{1,t}, X_{2,t}, \ldots, X_{k,t}, y_{t-1}) \tag{S5}$$

where $X_{i,t}$ denotes the environmental predictors and $y_{t-1}$ is abundance in the previous year.

Rather than modelling abundance as a linear function of these predictors, the input vector was projected into a high-dimensional dynamical state space through a fixed recurrent reservoir. The reservoir states evolved according to

$$\mathbf{r}_t = (1-\lambda)\,\mathbf{r}_{t-1} + \lambda \tanh\left(\mathbf{W}_{\text{res}}\mathbf{r}_{t-1} + \mathbf{W}_{\text{in}}\mathbf{u}_t\right) \tag{S6}$$

where $\mathbf{r}_t$ is the reservoir state vector, $\mathbf{W}_{\text{res}}$ and $\mathbf{W}_{\text{in}}$ are randomly initialized recurrent and input weight matrices, respectively, and $\lambda$ is the leaking rate controlling the reservoir memory. The recurrent weights remained fixed throughout training, with only the output layer being estimated from the data.

The expected abundance was then obtained by fitting a generalized linear model to the reservoir states,

$$\log(\hat{y}_t) = \beta_0 + \sum_{j=1}^{N_r} \beta_j r_{j,t}, \tag{S7}$$

where $\hat{y}_t$ represents the conditional expectation of the response, linked to the reservoir states $r_{j,t}$ through a log-linear relationship and $N_r$ is the reservoir size. As in the environmental GLM, a Tweedie response distribution with logarithmic link was used for the readout layer.

Both the Full and Basic Reservoir models were implemented using reservoirs composed of 2,000 recurrent units (neurons) with hyperbolic tangent activation functions. The Basic Reservoir differed only in replacing the environmental covariates with a constant input, such that the reservoir dynamics were driven exclusively by lagged abundance observations. We used one-year lagged abundance as the autoregressive input and fitted the readout layer using the same Tweedie generalized linear model described in Appendix S2. Predictive uncertainty was approximated by averaging over an ensemble of 500 independently initialized reservoirs with randomly perturbed dynamical hyperparameters, as described in the main text.

### S5 Forecast reliability analysis

Although forecast displacement ($D$) and uncertainty expansion ($U$) were selected as simple and interpretable diagnostics, they represent only two aspects of the predictive distribution. To evaluate whether these diagnostics captured the most informative properties of the predictive distribution, we additionally quantified a broader set of distributional descriptors, including measures of predictive spread, tail behaviour, asymmetry, entropy and multimodality. We also quantified a set of descriptors characterizing the historical training time series. We then compared both groups of descriptors between forecasts for which the Full Reservoir outperformed or underperformed AR(1). Differences between groups were summarized using Cliff's $\delta$ as the primary measure of effect size [5]. Two-sided Mann–Whitney tests [6] were performed to obtain Benjamini–Hochberg-adjusted $P$-values [7], which are reported in Supplementary Tables S2 and S3.

Measures of predictive spread, including forecast standard deviation, interquartile range and prediction interval widths, consistently exhibited the largest effect sizes (Cliff's $\delta \approx -0.35$), indicating that forecasts with broader predictive distributions were substantially more likely to be outperformed by the autoregressive benchmark (Table S2). By contrast, descriptors of tail behaviour, asymmetry, entropy and multimodality showed only weak associations with relative forecast skill (all $|\delta| < 0.15$).

Historical characteristics of the training time series were substantially less informative than descriptors derived from the predictive distribution. Whereas measures of predictive spread exhibited moderate effect sizes (Cliff's $|\delta| \approx 0.35$), all historical descriptors showed only negligible to small effects (maximum $|\delta| = 0.17$ for the coefficient of variation; Table S3). Together, these results indicate that forecast reliability is better anticipated from characteristics of the predictive distribution than from conventional descriptors of historical population dynamics.

**Table S2. Predictive distribution characteristics associated with relative forecast skill.** Comparison of distributional characteristics between forecasts for which the Full Reservoir outperformed AR(1) and forecasts for which it underperformed AR(1) at the first forecast horizon. Positive Cliff's $\delta$ values indicate larger values in forecasts for which the Full Reservoir achieved a lower (better) CRPS than AR(1), whereas negative values indicate larger values in forecasts for which the Full Reservoir achieved a higher (worse) CRPS than AR(1) [5]. P-values were adjusted using the Benjamini–Hochberg false discovery rate procedure [7].

| Category | Diagnostic | Cliff's $\delta$ | Effect size | FDR-adjusted $p$ |
|---|---|---|---|---|
| *Primary reliability diagnostics* | | | | |
| | Uncertainty expansion ($U$) | −0.393 | Medium | $1.72 \times 10^{-50}$ |
| | Forecast displacement ($D$) | −0.361 | Medium | $1.72 \times 10^{-50}$ |
| *Predictive spread* | | | | |
| | Forecast standard deviation | −0.356 | Medium | $2.92 \times 10^{-49}$ |
| | 90% prediction interval width | −0.355 | Medium | $3.17 \times 10^{-49}$ |
| | 98% prediction interval width | −0.354 | Medium | $4.94 \times 10^{-49}$ |
| | Interquartile range | −0.353 | Medium | $1.05 \times 10^{-48}$ |
| | Forecast coefficient of variation | 0.164 | Small | $1.25 \times 10^{-11}$ |
| *Tail behaviour* | | | | |
| | Outlier rate (IQR criterion) | 0.175 | Small | $5.92 \times 10^{-13}$ |
| | 98% PI / 50% PI ratio | 0.172 | Small | $1.39 \times 10^{-12}$ |
| | 90% PI / 50% PI ratio | 0.157 | Small | $6.70 \times 10^{-11}$ |
| | Excess kurtosis | 0.123 | Negligible | $1.23 \times 10^{-6}$ |
| *Asymmetry* | | | | |
| | Quantile skewness | 0.149 | Small | $5.41 \times 10^{-10}$ |
| | Skewness | 0.127 | Negligible | $6.13 \times 10^{-7}$ |
| *Entropy / complexity* | | | | |
| | Normalized histogram entropy | −0.173 | Small | $1.02 \times 10^{-12}$ |
| *Multimodality / normality* | | | | |
| | Number of KDE modes | −0.127 | Negligible | $1.04 \times 10^{-9}$ |
| | Normality test ($p$-value) | −0.127 | Negligible | $6.42 \times 10^{-7}$ |
| *Robust concentration* | | | | |
| | MAD / SD ratio | −0.168 | Small | $3.78 \times 10^{-12}$ |

**Table S3. Historical time-series characteristics associated with relative forecast skill.** Comparison of characteristics of the training time series between forecasts for which the Full Reservoir outperformed AR(1) and forecasts for which it underperformed AR(1) at the first forecast horizon. Positive Cliff's $\delta$ values indicate larger values in forecasts for which the Full Reservoir achieved a lower (better) CRPS than AR(1), whereas negative values indicate larger values in forecasts for which the Full Reservoir achieved a higher (worse) CRPS than AR(1) [5]. P-values were adjusted using the Benjamini–Hochberg false discovery rate procedure [7].

| **Training-series characteristic** | **Cliff's $\delta$** | **Effect size** | **FDR-adjusted $p$** |
|---|---|---|---|
| Coefficient of variation | 0.170 | Small | $7.16 \times 10^{-12}$ |
| Mean abundance | −0.141 | Negligible | $9.51 \times 10^{-9}$ |
| Variance | −0.116 | Negligible | $2.18 \times 10^{-6}$ |
| Lag-1 autocorrelation | −0.053 | Negligible | $3.86 \times 10^{-2}$ |
| Sample entropy | −0.052 | Negligible | $1.21 \times 10^{-1}$ |
| Training-series length | −0.033 | Negligible | $1.51 \times 10^{-1}$ |
